# Delineating memory reactivation in sleep with verbal and non-verbal retrieval cues

**DOI:** 10.1101/2023.03.02.530762

**Authors:** Anna á V. Guttesen, M. Gareth Gaskell, Scott A. Cairney

## Abstract

Sleep supports memory consolidation via the reactivation of newly formed memory traces. One way to investigate memory reactivation in sleep is by exposing the sleeping brain to auditory retrieval cues; a paradigm known as targeted memory reactivation (TMR). To what extent to acoustic properties of memory cues influence the effectiveness of TMR, however, has received limited attention. We addressed this question by exploring how verbal and non-verbal memory cues affect oscillatory activity linked to memory reactivation in slow-wave sleep. Fifty-one healthy adult males learned to associate visual stimuli with spoken words (verbal cues) and environmental sounds (non-verbal cues). Subsets of the verbal and non-verbal cues were then replayed during sleep, alongside previously unheard control cues. For a subset of the participants, the voice of the verbal cues was mismatched between sleep and learning. Memory cues (relative to control cues) prompted an increase in theta/alpha and spindle power, which have been heavily implicated in sleep-associated memory processing. Moreover, verbal memory cues were associated with a stronger increase in spindle power than non-verbal memory cues. There were no significant differences between the matched and mismatched conditions when analysing verbal memory cues in isolation. Our findings suggest that verbal memory cues may be more effective than non-verbal memory cues for triggering memory reactivation in sleep, as indicated by an amplified spindle response.

## Introduction

Sleep facilitates memory consolidation; the process through which information is retained in long-term memory. Sleep-associated memory gains were initially thought to arise from a passive protective mechanism, whereby sleep shields newly acquired memories from the interference posed by wakefulness (Jenkins & Dallenbach, 1924). However, more recent work has suggested that sleep also plays an active role in offline memory processing (Born & Wilhelm, 2012; Klinzing et al., 2019; Rasch & Born, 2013), such that hippocampus-dependent memories are repeatedly reactivated and gradually integrated with pre-existing representations in neocortex.

According to this *Active Systems Consolidation* framework, memory processing in sleep relies on finely-tuned interactions between the cardinal neural oscillations of slow-wave sleep (SWS): <1 Hz neocortical slow oscillations (SOs), 11-16 Hz thalamocortical sleep spindles and ∼80-100 Hz hippocampal ripples (Born & Wilhelm, 2012; Klinzing et al., 2019; Rasch & Born, 2013). Embedded within SOs, sleep spindles are thought to cluster reactivated memory units in the form of ripples to coordinate their transfer from hippocampus to neocortex for long-term storage (Staresina et al., 2015). Several studies in humans have provided compelling support for this view, demonstrating that patterns of brain activity observed at learning re-emerge during spindles, highlighting spindles as a candidate neural marker of memory reactivation in sleep (Bergmann et al., 2012; Schreiner et al., 2021; Schönauer et al., 2017).

Alongside spindles, SOs and ripples, the 4-8 Hz theta rhythm has also been implicated in overnight memory consolidation (Schreiner et al., 2018). Adaptations of the Active Systems framework suggest that spindle and theta oscillations work in unison to support memory reactivation and stabilisation during SWS, with theta activity representing the initial reactivation of newly formed memories and spindles signifying their subsequent reprocessing and migration to neocortex (Antony et al., 2019; Schreiner & Rasch, 2017).

Our understanding of sleep’s role in offline memory processing has been heavily influenced by the development of a memory cueing paradigm known as targeted memory reactivation (TMR; Rudoy et al., 2009). In a typical TMR study, participants form new memories that are associated with sounds at the time of learning. A subset of these sounds is then replayed during SWS to trigger the reactivation of their associated memory traces. A wide range of studies have shown that retention over sleep is improved for memories that are cued by TMR relative to those that are not (Antony et al., 2018; Cairney et al., 2016; Göldi et al., 2019; Schechtman et al., 2021; Schreiner, Lehmann, et al., 2015; Schreiner & Rasch, 2015; Schönauer et al., 2014), providing causal evidence that memory reactivation is a central mechanism of sleep-associated consolidation.

Beyond behavioural manifestations of memory reactivation, TMR has provided important evidence for the role of sleep spindles and theta oscillations in overnight consolidation. In humans, memory cues delivered during SWS trigger a transient increase in spindle activity, with the magnitude of this increase predicting later memory performance (Antony et al., 2018; Cox et al., 2014; Farthouat et al., 2017; Groch et al., 2017; Laventure et al., 2018; Lehmann et al., 2016; Schechtman et al., 2021; Schreiner, Lehmann, et al., 2015; Wang et al., 2019). TMR-evoked increases in spindle activity are often preceded by a surge in theta activity, consistent with the view that theta and spindle rhythms play complementary roles in offline memory processing (Bar et al., 2020; Farthouat et al., 2017; Groch et al., 2017; Joensen et al., 2022; Laventure et al., 2018; Lehmann et al., 2016; Oyarzún et al., 2017; Schechtman et al., 2021; Schreiner, Lehmann, et al., 2015; Schreiner & Rasch, 2015).

Although the majority of TMR studies have used environmental sounds (e.g. a dog barking) as memory cues (Antony et al., 2018; Cairney et al., 2016; Rudoy et al., 2009; Schechtman et al., 2021; Schönauer et al., 2014; Van Dongen et al., 2012; Vargas et al., 2019), several others have delivered verbal stimuli (i.e. spoken words) during sleep (Farthouat et al., 2017; Göldi et al., 2019; Lehmann et al., 2016; Schreiner, Göldi, et al., 2015; Schreiner & Rasch, 2015). Whether the effectiveness of TMR is any greater for verbal or non-verbal cues has received only limited attention, but is nevertheless an important question: by determining which type of memory cue engenders the greatest impact on offline memory processing, we can optimise TMR protocols and strengthen their potential utility as tools in education and healthcare (van der Heijden et al., 2022).

In previous work, we compared the impacts of verbal and non-verbal memory cues on sleep-associated consolidation. Participants associated visual stimuli with spoken words (verbal cues) or environmental sounds (non-verbal cues) at learning, which were then replayed during SWS (Cairney et al., 2017). Although cueing in SWS improved memory retention, the magnitude of this improvement was highly comparable across the verbal and non-verbal cueing conditions, suggesting that cue type had no impact on the memory enhancing effects of TMR.

However, the coarse behavioural measures used in our prior study might have been inadequate to detect differences in the effectiveness of verbal and non-verbal TMR cues. Given the recent evidence that spindle activity in sleep provides a neural marker of offline memory replay, sleep spindles (and other neural oscillations linked to memory reactivation in SWS) might provide a more reliable means of indexing the sleeping brain’s responsiveness to different types of memory cue.

Our earlier work also compared the memory effects of verbal TMR when the cues were acoustically matched or mismatched to those encountered at learning. Whereas matched verbal cueing led to a selective improvement in retention for the cued (but not the non-cued) associations, mismatched verbal cueing prompted a performance gain for both the cued and non-cued memories. This suggests that mismatched cueing might promote a generalised form of reactivation across a single learning context, and is in keeping with recent evidence that multiple memories can be simultaneously reactivated in response to a single TMR cue (Schechtman et al., 2022). However, to what extent the varied influences of matched and mismatched TMR cues are driven by disparate neural mechanisms cannot be inferred from our behavioural findings. The brain rhythms implicated in TMR might thus offer a solution to this outstanding question.

We report on a secondary analysis of sleep EEG data acquired in our previous study (Cairney et al., 2017) comparing the effects of TMR with verbal versus non-verbal memory cues (Experiment 1), and acoustically matched versus mismatched verbal memory cues (Experiment 2). Because our TMR protocol included verbal and non-verbal control stimuli that were not presented at learning, we could isolate the time-frequency responses associated with memory reactivation in sleep (i.e. memory cues > control cues). We exploited these time-frequency representations in our well-powered sample (N=51) to investigate whether the sleeping brain is more responsive to verbal or non-verbal memory cues (i.e. based on the evoked spindle/theta response), and assess whether the divergent behavioural influences of matched and mismatched cueing observed in our prior work are accompanied by distinct neural activity profiles.

## Materials and Methods

### Participants

Data from 51 healthy males were analysed (Experiment 1: N = 28, mean ± SD age = 20.32 ± 1.54, Experiment 2: N = 23, mean ± SD age = 20.96 ± 2.38). Screening questionnaires indicated that participants had no history of sleep, neurological or psychiatric disorders, were non-smokers and were not using any psychoactive medications. Following standard practices in our lab, participants were instructed to refrain from alcohol and caffeine for 24 hours before the start of the study (Ashton et al., 2019; Cairney, Lindsay, et al., 2018; Guttesen et al., 2022; Strachan et al., 2020). The Pittsburgh Sleep Quality Index (Buysse et al., 1989) indicated that all participants had a normal pattern of sleep in the month preceding the study. Participants provided written and informed consent and the study was approved by the Research Ethics Committee of the Department of Psychology at the University of York.

### Procedure

#### Experiment 1

##### Evening

Figure 1 illustrates the experimental procedure. Participants arrived at the sleep laboratory at 9:30pm (± 30 min) and were wired up for sleep EEG monitoring. They were informed that the study was about the role of sleep in memory consolidation but were not told about the TMR manipulation. Training comprised a paired-associates task and included a learning phase and a test phase. At learning, participants associated each of 56 visually presented words with an auditory stimulus (28 verbal stimuli and 28 non-verbal stimuli). All verbal cues were presented in a male or female voice (counterbalanced across participants). At test, participants were presented with each of the auditory stimuli and instructed to type the associated words. If participants failed to correctly recall >60% of the target words, the training (learning and test) was repeated. Participants then completed a final pre-sleep test where they were assessed on all 56 paired associates (following the same procedures as the prior test phases). See Cairney et al. (2017) for a more detailed account of the experimental task and associated procedures.

**Figure 1:**
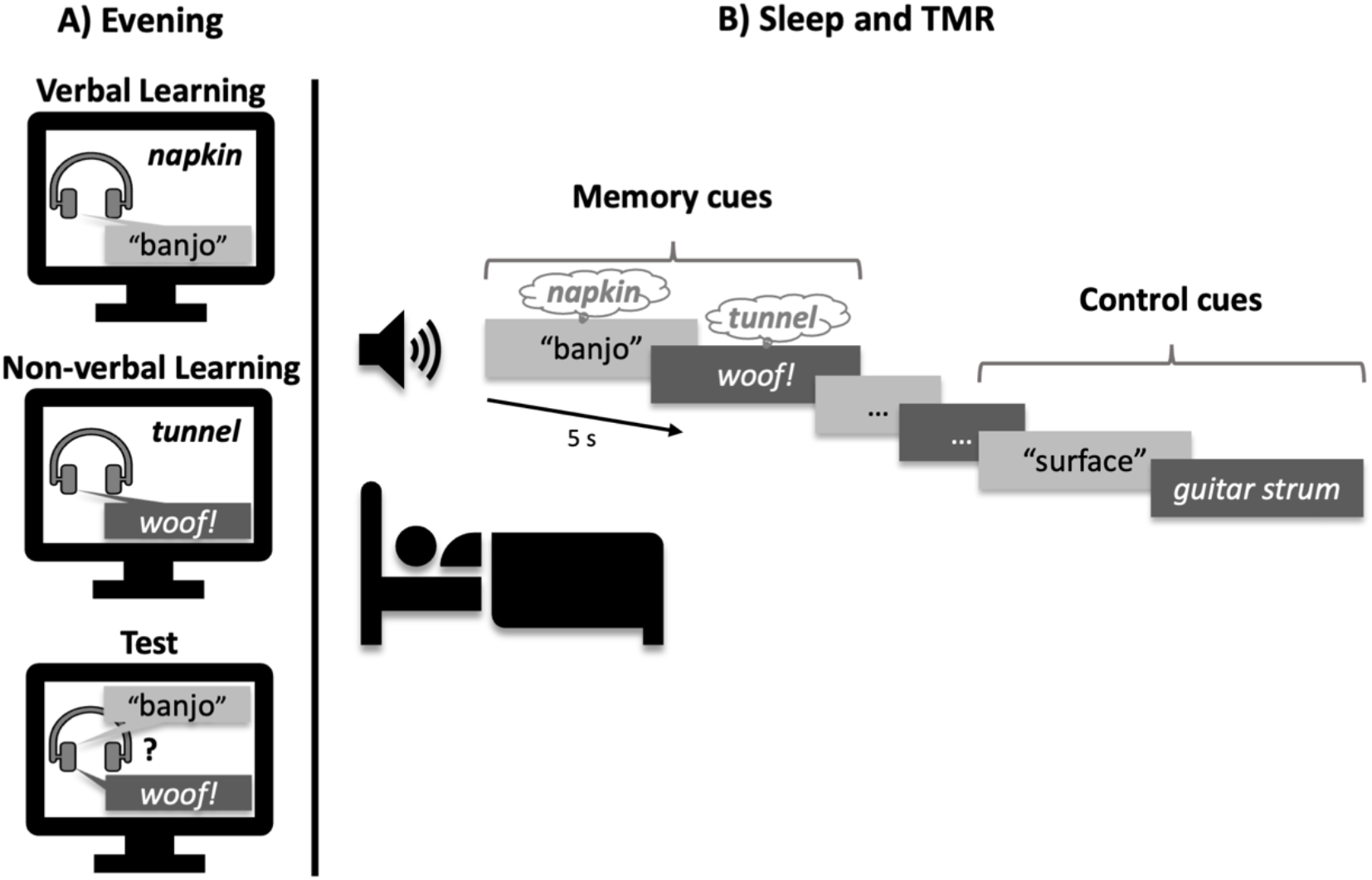
Experimental procedure. **A)** Participants learned to associate visually presented words with verbal and non-verbal auditory cues. **B)** Verbal (light-grey) and non-verbal (dark-grey) auditory cues were replayed during slow-wave sleep (order randomised). Previously unheard control cues were also played. Verbal cues were presented in the same voice as at learning (matched) in Experiment 1 and in a different voice to learning in Experiment 2 (mismatched). Non-verbal TMR cues were identical to those heard at learning in both experiments. TMR = targeted memory reactivation.

##### TMR cues

For the paired associates scored as correct on the final pre-sleep test, half of the verbal cues and half of the non-verbal cues were randomly selected and intermixed for replay in SWS. Two additional control cues that were not present at learning (the spoken word “surface” and the sound of a guitar strum) were randomly interspersed within the memory cues. These were played the same number of times that their corresponding verbal and nonverbal cues were replayed, and at the beginning of the TMR rounds (2 verbal and 2 non-verbal) to ensure that participants’ sleep would not be disturbed during auditory stimulation.

##### Overnight sleep

Lights were turned out at ∼11 pm. White noise was played throughout the night to habituate participants to auditory stimulation (39 dB). TMR began after participants had exhibited at least 2 min of continuous SWS (as determined via online sleep EEG monitoring). Memory cues were played at 5 s intervals and white noise intensity was lowered during the replay of each cue to promote acoustic clarity. Because the number of memory cues varied across participants, null events (i.e. events with no stimulation) were randomly intermixed between the cues so that each round of TMR always lasted 290 s. TMR was repeated throughout the first two cycles of SWS with 1-min intervals placed between each completed round. Cueing was stopped if participants transitioned from SWS to another sleep stage or wakefulness, or showed signs of microarousals, but was restarted if they returned to SWS. At ∼7 am, participants were woken up, unless they were in SWS or REM, in which case they were allowed to continue sleeping until they woke up or reached N1 or N2.

### Experiment 2

Experiment 2 followed the same procedures as Experiment 1, with the exception that the verbal cues were presented in a male voice at training and test, and in a female voice during sleep. The non-verbal cues were identical to those of Experiment 1 and did not differ between learning and TMR.

### Stimuli

#### Verbal cues

Thirty-five monosyllabic and disyllabic words (mean ± SD syllable count = 1.54 ± 0.51) were taken from the University of South Florida (USF) word association, rhyme, and word fragment norms (Maki et al., 2004; Nelson et al., 1998) for use as verbal cues. The words were recorded using two separate speakers; one male and one female. The male and female word recordings were of similar duration (mean ± SD ms: male = 769.29 ± 104.95, female = 774.80 ± 99.14, t(34) = 0.49; p = .63). An additional word (“surface”) was taken from the USF norms for use as a spoken control cue (male duration = 990 ms; female duration = 950 ms). The abstract nature of this control word was intentional so that it remained distinct from the verbal memory cues.

#### Non-verbal cues

Thirty-five environmental sounds were taken from previous studies (Oudiette & Paller, 2013; Rudoy et al., 2009) and from freesound.org for use as non-verbal cues. The sounds were similar in length to both the male and female versions of the verbal cues (mean duration ± SD ms = 740.97 ± 156.29, F(2,102) = 0.76; p = .47). Additionally, the sound of a guitar strum (524 ms) was taken from Rudoy et al. (2009) to serve as the non-verbal control cue.

#### Visual stimuli

A further seventy monosyllabic and disyllabic words were taken from the USF norms for use as visual targets in the in the verbal and non-verbal paired associates. Each word was paired with a verbal and non-verbal auditory cue, resulting in two 35-item sets of verbal paired associates (verbal A and B) and two 35-item sets of non-verbal paired associates (non-verbal A and B). For the experiments, if the verbal paired associates were taken from set A, the non-verbal paired associates were taken from set B and vice-versa (counterbalanced across participants). None of the paired associates had a clear semantic link.

### Equipment

#### Sleep EEG

Sleep was monitored using an Embla N7000 PSG system with RemLogic version 3.4 software. Gold-plated electrodes were attached to the scalp in accordance with the international 10-20 system at frontal (F3 and F4), central (C3 and C4) and occipital (O1 and O2) locations, and were each referenced to the contralateral mastoid (M1 and M2). Left and right electrooculography electrodes were attached, as were electromyography electrodes at the mentalis and submentalis bilaterally, and a ground electrode was attached to the forehead. Each electrode had a connection impedance of < 5 kΩ and all online signals were digitally sampled at 200 Hz. Sleep scoring was conducted on the referenced central electrodes (C3 and C4) in accordance with standardised criteria (Iber, 2007).

#### TMR

TMR was implemented with Presentation v17.0 (Neurobehavioral Systems, Inc.). Auditory cues were played via a speaker placed ∼1.5m above the bed, which was connected to an amplifier in a separate control room.

### EEG analyses

All data preprocessing and analyses were conducted in MATLAB version 2019a using FieldTrip toolbox version 10/04/18 (Oostenveld et al., 2011).

#### Preprocessing

Sleep EEG data were re-referenced to the linked mastoids (average of M1 and M2), notch filtered at 49-51 Hz, high-pass filtered at 0.5 Hz and then segmented into trials (−1s to 3.5 s around cue onset). Using FieldTrip’s Databrowser function, the data were first visually inspected for noisy channels (none were identified). Automatic artifact rejection was then implemented using FieldTrip’s automated artifact rejection function (*ft_artifact_zvalue*). In this step, muscle artifacts at 15-32 Hz (Brunner et al., 1996) were exaggerated using filters and z-transformations (0.1 s padding on each side of the artifact) and then removed (mean ± SD trials rejected across all participants in both experiments = 3.96 ± 2.26). The remaining artifacts were manually rejected based on visual inspection via FieldTrip’s Databrowser (mean ± SD noisy trials rejected across all participants in both experiments = 4.14 ± 5.23). Trials that fell outside of sleep stages N2 or SWS were excluded prior to analysis (mean ± SD trials removed across all participants in both experiments = 7.55 ± 10.84). Table 1 shows the number of trials in each condition after artifact rejection and trial removal.

**Table 1:** TMR trials per condition. Control cues were intermixed with the memory cues and also played at the beginning of each TMR set to ensure that participants’ sleep would not be disturbed during auditory stimulation (hence a higher number of control cues than memory cues). Verbal memory and control cues were presented in the same voice as at learning (matched) in Experiment 1 and in a different voice to learning in Experiment 2 (mismatched). Data are shown as mean ± SD.

|  | Memory cues |  | Control cues |  |
| --- | --- | --- | --- | --- |
|  | Experiment 1 | Experiment 2 | Experiment 1 | Experiment 2 |
| **Verbal** | *Matched*<br>$89.68 \pm 35.60$ | *Mismatched*<br>$78.22 \pm 29.37$ | *Matched*<br>$108.50 \pm 41.81$ | *Mismatched*<br>$97.17 \pm 33.82$ |
| **Non-verbal** | $88.57 \pm 38.85$ | $79.57 \pm 32.18$ | $107.64 \pm 46.83$ | $97.70 \pm 37.45$ |
| **Total** | $169.02 \pm 66.58$ | | $206.55 \pm 78.93$ | |

#### Time-frequency analyses

Time-frequency representations (TFRs) were calculated for frequencies ranging from 4-30 Hz. Data were convolved with a 5-cycle Hanning taper in 0.5 Hz frequency steps and 5 ms time steps using an adaptive window-length (i.e. where window length decreases with increasing frequency, e.g. 1.25 s at 4 Hz, 1 s at 5 Hz etc). TFRs were converted into % power change relative to a -300 to -100 ms pre-cue baseline window. This window was chosen to mitigate baseline contamination by post-stimulus activity while preserving proximity to cue onset (Cairney, Guttesen, et al., 2018).

#### Event-related potentials

For event-related potentials (ERPs), data were high-pass filtered at 0.5 Hz and low-pass filtered at 30 Hz. Data were baseline-corrected relative to a -200 ms to 0 ms pre-cue window (Cairney, Guttesen, et al., 2018).

#### Statistics

ERP and TFR analyses were corrected for multiple comparisons using FieldTrip’s non-parametric cluster-based permutation method with 1000 randomisations (increased to 1500 when the standard deviation of the p-value crossed the alpha-value, Meyer et al., 2021). All time-frequency clusters were defined by channel * time * frequency (4-30 Hz, cluster threshold p < .05, two-tailed). The time window of interest in the TFR was 0.3-2.5 s (Cairney, Guttesen, et al., 2018). ERP clusters were defined by time (averaged across channels) and based on a 0-2.5 s time window of interest (4-30 Hz, cluster threshold p < .05, two-tailed).

Because Experiments 1 and 2 were highly similar (with the only difference being the match vs mismatch of speaker between training and verbal TMR), we first collapsed the data across experiments and compared memory cues to control cues with a dependent-samples analysis. To assess the effects of cue type (verbal vs non-verbal) on oscillatory activity, we used a factorial approach. We calculated the grand average difference for the contrast [memory cues > control cues] within each condition (verbal cues and non-verbal cues), and then entered these contrasts into a dependent-samples analysis (verbal^memory cues>control cues^ > non-verbal^memory cues>control cues^). We also used a factorial approach to assess the effects of matched vs mismatched verbal cues (i.e. matched^memory^ cues>control cues> mismatched^memory cues>control cues^). However, because the matched and mismatched conditions were collected across Experiment 1 and Experiment 2, respectively, we used an independent-samples comparison in this analysis. Cohen’s dz effect sizes were based on the largest identified clusters by averaging power across the time points, frequencies and channels that contributed to the clusters at any point.

## Results

### Memory cues and control cues

First, we examined the sleeping brain’s response to memory cues memory cues (Figure 2a) and control cues (Figure 2b). Significant differences in the time-frequency representation (TFR) were observed for memory cues (vs control cues, p < .05, Figure 2c). The identified clusters showed an increase in theta/alpha power (∼4-11.5 Hz) across both hemispheres at ∼0.3-0.9 s (left: d_z_ = .52, right: d_z_ = .48), which was followed by an increase in spindle/beta power (∼10.5-20 Hz) at ∼0.8-1.7 s (left: d_z_ = .51, right: d_z_ = .56). There was also a later decrease in power across a wider spindle/beta band (∼12-26 Hz) in both hemispheres (∼1.8-2.5 s, left: d_z_ = -.38, right: d_z_ = -.46, Figure 2c and 2d). To distinguish between the early increase (∼10.5-20 Hz) and later decrease in activity across the spindle/beta bands (∼12-26 Hz), we refer to the former as spindle power and the latter as spindle/beta power hereafter (because the latter encompasses a wider frequency range). However, it should be noted that our “spindle” band encompasses higher frequencies than would normally be considered spindle activity.

**Figure 2:**
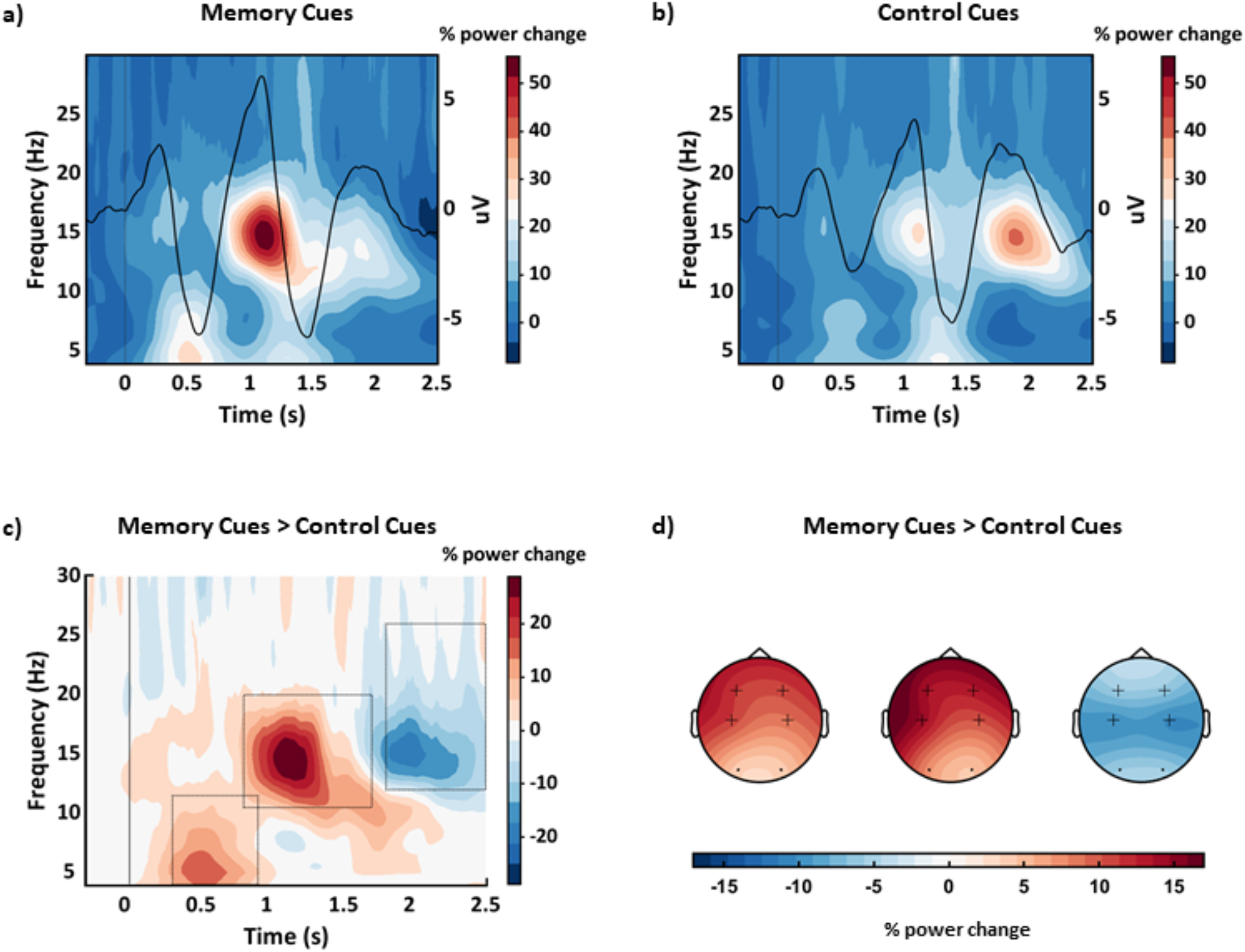
Memory cues and control cues. Grand average time-frequency representations with superimposed event-related potentials (baseline corrected and averaged across all channels) for **a)** memory cues and **b)** control cues. Memory cues > control cues in **c)** time-frequency representation and **d)** topographical representation. The rectangles in 2c illustrate timing and frequency windows of topographical distributions in 2d (left to right). Crosses represent the channels contributing to the largest clusters (i.e. F3, F4, C3 and C4). Data are collapsed across Experiments 1 and 2.

The ERP evoked by memory cues (Figure 2a) was also significantly stronger than that evoked by control cues (Figure 2b, p <.05), with three clusters at ∼0.4-0.7 s (negative, d_z_ = -.70), ∼0.9-1.3 s (positive, d_z_ = .59) and ∼1.4-1.8 s (negative, d_z_ = -.64).

### Verbal and non-verbal memory cues

Next, we examined whether verbal and non-verbal memory cues evoke distinct patterns of oscillatory activity during sleep. We subtracted the evoked control cue response (Figure 3b and 3d) from the memory cue response (Figure 3a and 3c), separately for verbal and non-verbal cues, leading to a 2×2 factorial design (verbal^memory cues>control cues^ > non-verbal^memory cues>control cues^). A significant difference emerged (p < .05), which corresponded to an increase in spindle activity (∼10.5-16.5 Hz) across the right hemisphere at ∼0.5-1 s (d_z_ = .27, Figure 3e and 3f). Interestingly, post-hoc tests of the identified cluster revealed a stronger spindle response for verbal memory cues relative to both non-verbal memory cues (p = .008) and verbal control cues (p < .001). No significant differences were observed between the non-verbal memory cues and non-verbal control cues (p = .800), nor between the verbal and non-verbal control cues (p = .129, all Bonferroni corrected).

**Figure 3:**
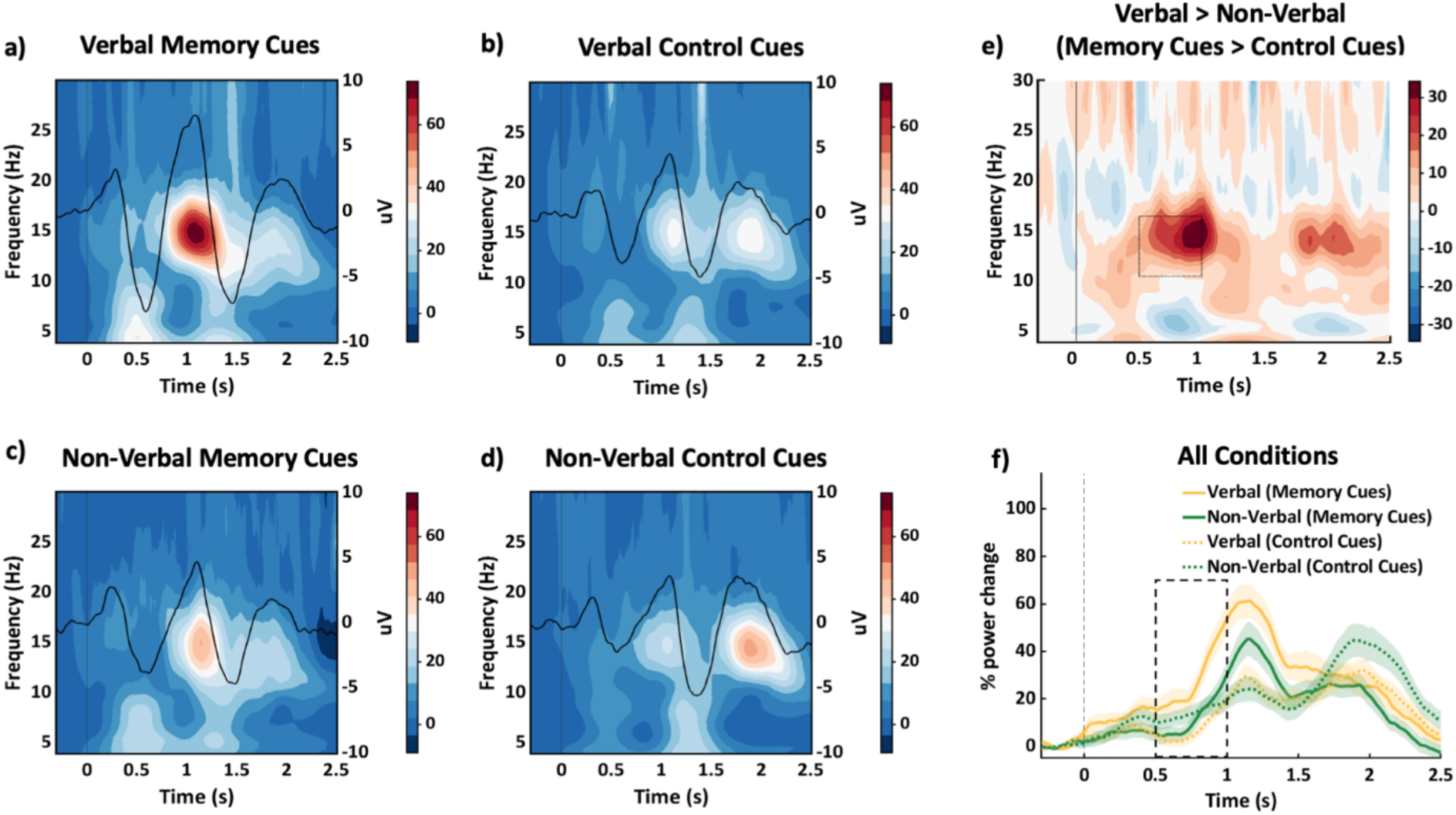
Verbal cues and non-verbal cues. Grand average time-frequency representations with superimposed event-related potentials (baseline corrected and averaged across all channels) for **a)** verbal memory cues, **b)** verbal control cues, **c)** non-verbal memory cues and **d)** non-verbal control cues. **e)** Grand average difference for the contrast: verbalmemory cues>control cues > non-verbalmemory cues>control cues. The rectangle illustrates the timing (∼0.5-1 s) and frequency (∼10.5-16.5 Hz) which contributed to the largest cluster. Colour bars represent % change. **f)** Mean (±SEM) power change over time for all four conditions (collapsed across the channels [F4 and C4] and frequencies [10.5-16.5 Hz] that contributed to the largest cluster in **e**). The rectangle illustrates the approximate timing which contributed to the cluster. Data are collapsed across Experiments 1 and 2.

We employed the same factorial approach to our ERP analysis and observed a significant difference (p < .05, verbal^memory cues>control cues^ > non-verbal^memory cues>control cues^), with two clusters at ∼0.8-1 s (positive, d_z_ = .47) and ∼1.2-1.5 s (negative, d_z_ = -.50).

### Matched and mismatched memory cues

Finally, we investigated whether verbal memory cues that are acoustically matched to those heard at learning evoke distinct neural responses to those arising from mismatched verbal cues (i.e. same vs different voice). We subtracted the evoked control cue response from the memory cue response, separately for the matched (Experiment 1) and the mismatched conditions (Experiment 2), leading to a 2×2 mixed factorial design (matched^memory cues > control cues^ > mismatched^memory cues > control cues^). However, no significant interaction emerged (Figure 4).

**Figure 4.**
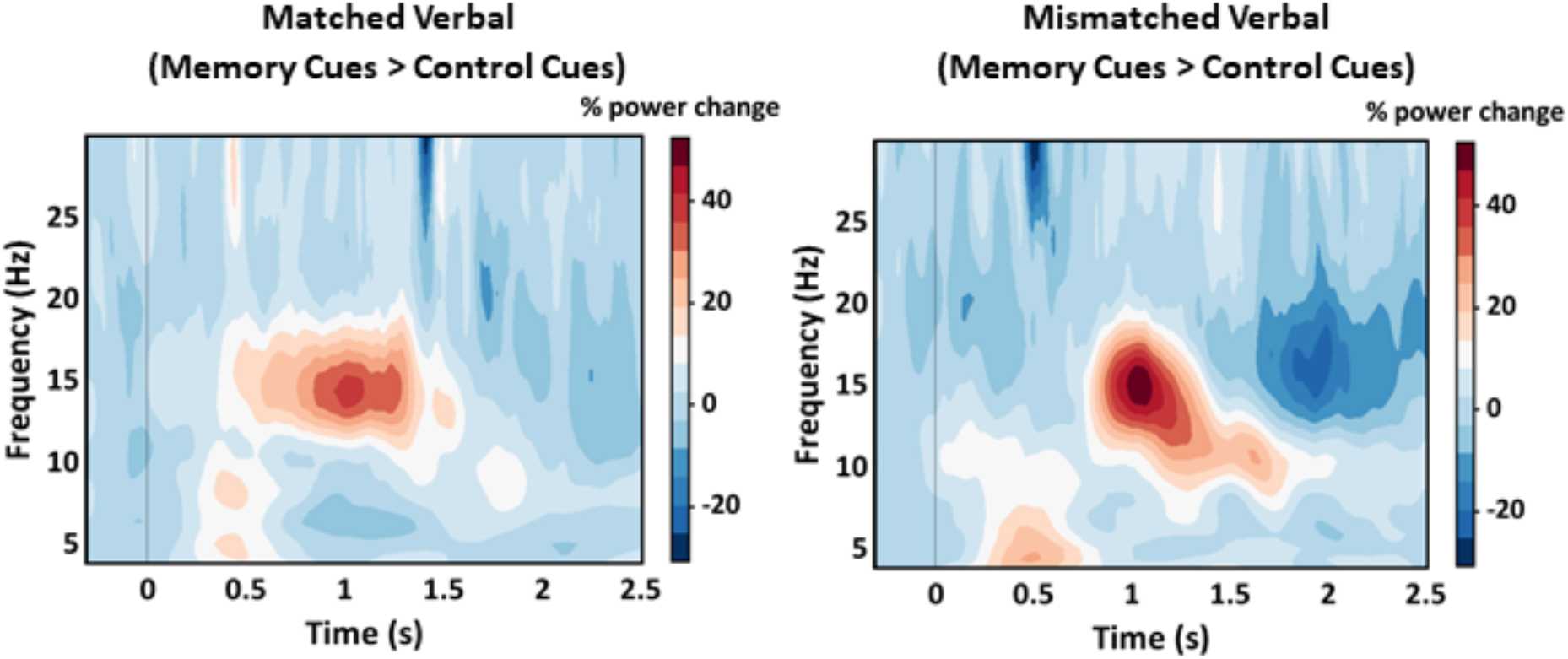
Matched and mismatched verbal cues. Grand average time-frequency representations (memory cues > control cues) for the matched verbal cues (Experiment 1) and mismatched verbal cues (Experiment 2). No significant effects were observed (p > .05).

The same factorial approach was applied to our ERP analysis, but again no significant interaction emerged (verbal^memory cues>control cues^ > non-verbal^memory cues>control cues^).

## Discussion

Overnight memory consolidation is achieved through the reactivation of newly formed memory traces in SWS (Born & Wilhelm, 2012; Klinzing et al., 2019; Rasch & Born, 2013). Studies employing the targeted memory reactivation (TMR) paradigm have provided crucial support for this hypothesis, as well as the view that sleeping brain rhythms, including theta and spindle oscillations, play a central role in overnight memory processing (Antony et al., 2019; Schreiner & Rasch, 2017). In the current study, we explored whether neural activity evoked by TMR in SWS is influenced by: 1) the acoustic properties of memory cues, and 2) the acoustic overlap between cues presented at learning and sleep.

Relative to previously unheard control cues, memory cues prompted an initial increase in theta/alpha and then spindle power, which was followed by a decrease in spindle/beta power. Importantly, the evoked spindle response was significantly stronger for verbal cues than non-verbal memory cues, suggesting that verbal auditory stimuli might be particularly potent triggers of memory reactivation in sleep. There were no significant differences in the evoked response to verbal cues that were acoustically matched or mismatched to learning.

The observed increase in theta/alpha and then spindle power for memory cues relative to control cues is in keeping with a number of prior studies that examined the neural correlates of TMR (Bar et al., 2020; Cairney, Guttesen, et al., 2018; Groch et al., 2017; Göldi et al., 2019; Laventure et al., 2018; Lehmann et al., 2016; Schechtman et al., 2021; Schreiner, Lehmann, et al., 2015). Based on these findings, it has been argued that theta and spindle rhythms play complementary roles in overnight consolidation, with the former supporting memory reinstatement and the latter facilitating memory strengthening and stabilisation (Schreiner & Rasch, 2017). These previous studies reporting an increase in delta/theta and spindle activity during TMR have used a wide variety of memory cues, including auditory (Cairney, Guttesen, et al., 2018; Groch et al., 2017; Göldi et al., 2019; Schechtman et al., 2021; Schreiner, Lehmann, et al., 2015) and olfactory stimuli (Bar et al., 2020; Laventure et al., 2018), and have assessed the retention of memories with varying emotional properties (Lehmann et al., 2016) across both declarative (Bar et al., 2020; Cairney, Guttesen, et al., 2018; Groch et al., 2017; Göldi et al., 2019; Lehmann et al., 2016; Schechtman et al., 2021; Schreiner, Lehmann, et al., 2015) and non-declarative domains (Laventure et al., 2018). Hence, spindle and theta oscillations appear to play a domain-general role in sleep-associated memory reactivation and consolidation.

The initial TMR-evoked increase in spindle activity was followed by a decrease in spindle/beta power for memory cues relative to control cues. This was a surprising finding which, to our knowledge, has not been reported previously. A late decrease in spindle/beta power might reflect a period of spindle refractoriness following an initial surge in spindle-mediated memory processing. This is in line with a recent theoretical framework that describes how spindle refractory periods protect memories from interference during critical reprocessing windows (Antony et al., 2019; Schreiner & Rasch, 2017). Somewhat relatedly, decreases in alpha/beta power have been linked to successful learning and retrieval during wakefulness (Griffiths et al., 2016; Griffiths, Martín-Buro, Staresina, & Hanslmayr, 2021; Hanslmayr et al., 2012; Hanslmayr et al., 2011), and are thought to reflect reinstatement of an optimal neurophysiological state for suppressing noise (Griffiths, MartÍn-Buro, Staresina, Hanslmayr, et al., 2021). Beta desynchronization during TMR might thus reflect engagement of a similar noise-reducing mechanism to facilitate overnight consolidation.

Intriguingly, the increase in spindle activity observed during TMR was amplified for verbal relative to non-verbal memory cues. Given the putative function of spindles in sleep-associated memory processing (Antony et al., 2019; Schreiner & Rasch, 2017), our data might suggest that verbal cues are more effective than non-verbal cues for triggering memory reactivation in sleep. Indeed, post-hoc tests revealed a significant increase in the evoked spindle response for verbal memory cues relative to verbal control cues (and non-verbal memory cues), but no such difference between verbal control cues and non-verbal control cues, indicating that our findings do not simply reflect generalised differences in the processing of environmental sounds and spoken words. Although the stimuli were matched for duration, it is worth noting that other aspects of the stimulus sets could explain this observed difference, such as their acoustic properties. However, based on current understanding of the phonological and semantic processes underpinning spoken word recognition during wakefulness (Gaskell & Mirkovic, 2016; McMurray et al., 2022), it is possible that the sleeping brain has better access to meaning when presented with verbal relative to non-verbal memory cues, with an enhanced spindle response reflecting engagement of multi-level decoding pathways.

We observed no significant differences in the evoked EEG response for verbal memory cues that were acoustically matched or mismatched to those presented at learning (based on the speaker’s voice). This is at odds with our behavioural data (Cairney et al., 2017), which show the typical selective benefit of TMR for cued memories in the matched condition, but a generalised benefit for all cued and non-cued memories in the mismatched condition. Recent work has shown that, although TMR delivered in SWS evokes an increase in delta/theta (0.5-8 Hz) and spindle (11-16 Hz) power irrespective of whether the memory cues are associated with one or multiple targets, the magnitude of this increase is modulated by the number of targets (Schechtman et al., 2021). Hence, differences in the evoked response to matched and mismatched cues might be observed as gradual increases in power (according to the number of potential targets reactivated by a mismatched cue). Because our matched and mismatched conditions comprised different groups (from Experiment 1 and Experiment 2, respectively), we were unable to address this possibility in the current dataset, but it remains an open question for future research. Overall, the EEG data reported here complement and extend the previously reported behavioural data: alongside the commonalities, both provide unique insights into how different types of stimuli can affect memory processing during sleep.

A potential limitation of our protocol is that it included only two control cues (one verbal and one non-verbal), which were repeatedly replayed and intermixed with a much larger number of distinct memory cues. Event-related potentials were thus significantly smaller for control cues than memory cues, reflecting habituation of the neural response that might have influenced our other findings (Megela & Teyler, 1979). Future research comparing neural signatures of memory cues and control cues should ensure that the control cues are optimally designed, by matching the two types of stimuli in terms of auditory and linguistic characteristics, as well as their frequency of repetition. Finally, given the exploratory nature of this secondary analysis, future studies are needed to confirm our findings with an a-priori hypothesis-driven approach.

In conclusion, we found that memory cues evoke increases in theta and spindle power, which have been previously linked to memory reinstatement and stabilization during sleep. We also showed, for the first time, that the TMR-evoked spindle response is stronger for verbal memory cues than non-verbal memory cues, suggesting that verbal stimuli are more effective triggers of memory reactivation in the sleeping brain. However, we did not observe any differences when comparing neural responses to verbal memory cues that were matched or mismatched to those encountered at learning. Taken together, these findings provide novel insights into how the sleeping brain processes memory cues with distinct acoustic properties.

## Acknowledgements

We are grateful to Simon Hanslmayr and Aidan Horner for fruitful discussions of the data.

